# Structural basis of SARS-CoV-2 polymerase inhibition by Favipiravir

**DOI:** 10.1101/2020.10.19.345470

**Authors:** Qi Peng, Ruchao Peng, Bin Yuan, Min Wang, Jingru Zhao, Lifeng Fu, Jianxun Qi, Yi Shi

## Abstract

The outbreak of severe acute respiratory syndrome coronavirus 2 (SARS-CoV-2) has developed into an unprecedented global pandemic. Nucleoside analogues, such as Remdesivir and Favipiravir, can serve as the first-line broad-spectrum antiviral drugs against the newly emerging viral diseases. Recent clinical trials of these two drugs for SARS-CoV-2 treatment revealed antiviral efficacies as well as side effects with different extents^1–4^. As a pyrazine derivative, Favipiravir could be incorporated into the viral RNA products by mimicking both adenine and guanine nucleotides, which may further lead to mutations in progeny RNA copies due to the non-conserved base-pairing capacity^5^. Here, we determined the cryo-EM structure of Favipiravir bound to the replicating polymerase complex of SARS-CoV-2 in the pre-catalytic state. This structure provides a missing snapshot for visualizing the catalysis dynamics of coronavirus polymerase, and reveals an unexpected base-pairing pattern between Favipiravir and pyrimidine residues which may explain its capacity for mimicking both adenine and guanine nucleotides. These findings shed lights on the mechanism of coronavirus polymerase catalysis and provide a rational basis for developing antiviral drugs to combat the SARS-CoV-2 pandemic.

## Main

*Coronaviridae* family includes many life-threatening human pathogens, such as SARS-CoV^6^, Middle East Respiratory Syndrome coronavirus (MERS-CoV)^7^ and the ongoing pandemic SARS-CoV-2^8,9^. Coronaviruses harbor a non-segmented positive-sense RNA genome with 5’-cap and 3’-polyA tail, which can directly serve as mRNA to guide the production of viral proteins^10^. A total of 4 structural proteins and at least 16 non-structural proteins (nsps) as well as 8 accessory proteins are encoded by the coronavirus genome. Among them, nsp12 is the core catalytic subunit of viral RNA-dependent RNA polymerase (RdRp) complex, which executes transcription and replication of the viral genomic RNA^11^. Two cofactor subunits, nsp7 and nsp8, associate with nsp12 to constitute an obligatory core polymerase complex to confer processivity for RNA elongation^12^. To achieve complete transcription and replication, a panel of other nsps are involved to accomplish other enzymatic functions, including the nsp14-nsp10 exonuclease for proofreading^13–15^, the nsp13 helicase for RNA unwinding^16–19^, the nsp14 N7-methyltransferase and the nsp16-nsp10 2’-O-methyltransferase for capping^20–24^. Due to the key roles of polymerase complex for viral replication, it has long been thought of as a promising antiviral drug target^25^. Recently, we and other groups have determined the structures of SARS-CoV-2 core polymerase complex in both apo and RNA-bound states^26–30^, providing important information for structure-based antiviral drug design.

In the efforts to combat SARS-CoV-2 pandemic, Remdesivir was initially expected as a highly competent drug candidate for disease treatment. However, the recently disclosed outcome of clinical trials revealed controversial efficacies as well as some cases of side effects^1,2^. The clinical improvement rate of patients receiving Remdesivir within 7 days was only 3% and ~66% subjects developed adverse effects^1,2^. In contrast, another broad-spectrum antiviral Favipiravir showed good promise that 61.21% patients were clinically recovered at day 7 of treatment and 31.90% subjects were reported with side effect manifestations^3,4^. Both of them are administered as pro-drugs and should be processed into triphosphate forms as nucleotide mimicries to interfere with RNA synthesis. The active form of Remdesivir has a similar base moiety to adenine nucleotide which can be incorporated into the growing strand of RNA product using an uracil nucleotide as the template^30^. Besides, the cyanogroup in the ribose ring may cause steric clash with polymerase residues during elongation, resulting in aberrant termination of RNA synthesis^29,31^. It is yet unclear how Favipiravir could be recognized by the polymerase and disturb the faithful process of RNA production.

To investigate the mechanism for the antiviral efficacy of Favipiravir, we performed in vitro primer-extension assays using a template derived from the 3’-untranslated region (3’-UTR) of the authentic viral genome. The +1 catalytic position was inserted with different template residues accordingly to allow only one nucleotide to be incorporated in the presence of each individual nucleotide triphosphate (NTP) substrate (Fig. 1). Even though the product strand was supposed to grow by only one nucleotide, some larger RNA products with two or three nucleotides extension were generated for each specific NTP substrate (Fig. 1c and d). This phenomenon suggests the SARS-CoV-2 polymerase is prone to mis-incorporate unwanted nucleotides into the product RNA, resulting in low fidelity for transferring the genomic information during transcription and replication. Similar observations were also reported recently that SARS-CoV-2 polymerase is more tolerant for mismatches between template and product residues than other viral RdRps^5^, which further highlights the requirement for the proofreading nuclease nsp14 to maintain the integrity of viral genome. Interestingly, Favipiravir could be incorporated into the RNA product with similar efficiencies to those of ATP or GTP substrates guided by U or C template residues, respectively (Fig. 1c). In contrast, Remdesivir could only be incorporated with a U in the template strand (Fig. 1d). These evidences demonstrate that Favipiravir is a universal mimicry for purine nucleotides, different from other specific nucleotide analogues, such as Remdesivir^29,30^ and Sofosbuvir^32^.

**Fig. 1.**
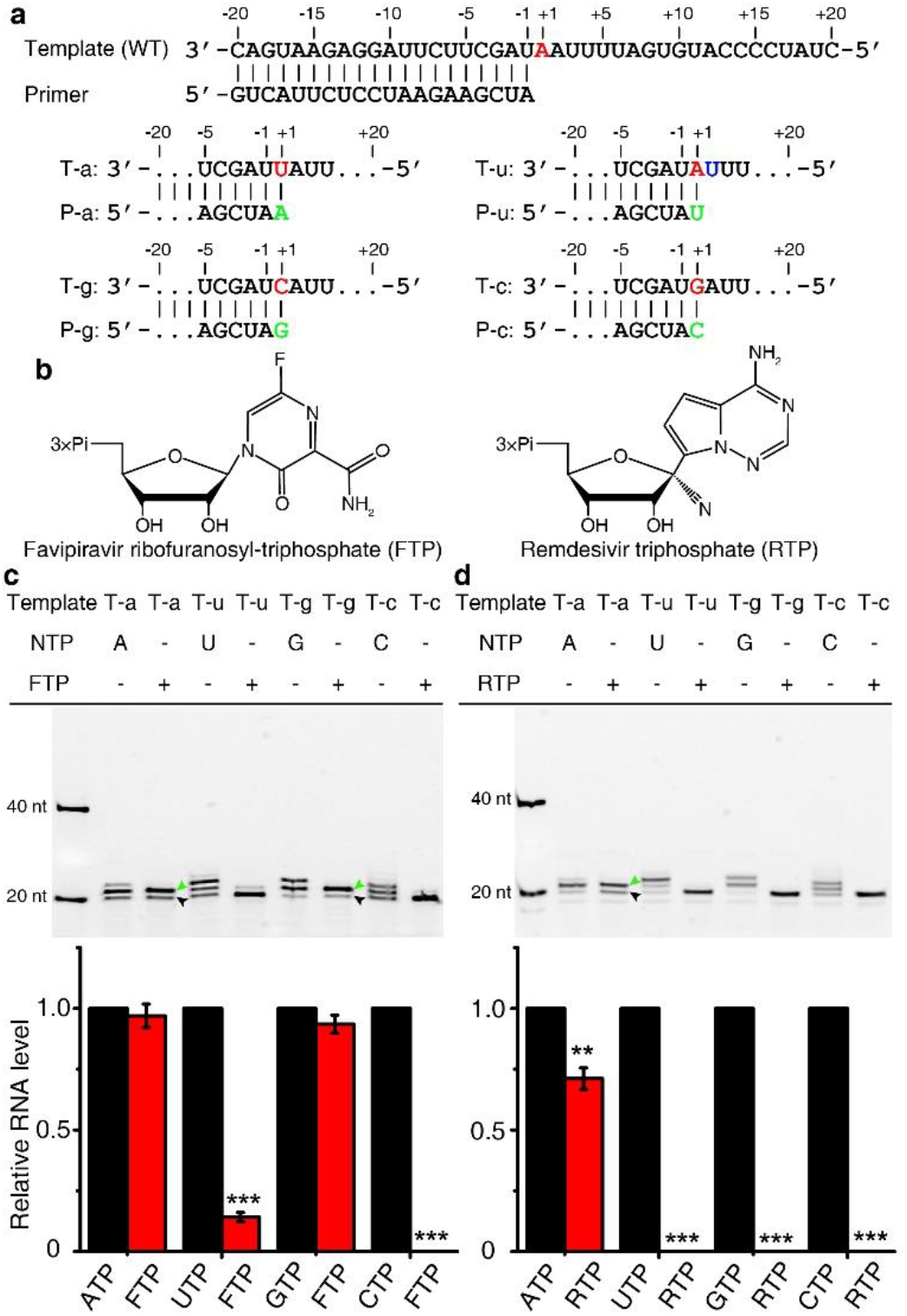
Incorporation of Favipiravir and Remdesivir into RNA products. **a,** RNA template and primer sequences engineered for single nucleotide extension. The +1 template residues are highlighted in red color and the nucleotide to grow is indicated by green. **b,** Structures of the metabolically active form of Favipiravir and Remdesivir. **c-d,** Single nucleotide extension assays for Favipiravir and Remdesivir. The remaining 20-nt primer and the 21-nt single nucleotide extension products are indicated by black and green arrowheads respectively. Different extents of extra-extension products are also generated in reactions for each individual NTP substrate, which may result from mis-matches between template residues and incoming nucleotides. The products were quantified by integrating the intensity of each band. The incorporation efficiency of Favipiravir for each specific template was compared to the corresponding NTP substrate. The differences were estimated by two-tailed student t-test with results from three independent experiments using different protein preparations. *, P<0.05; **, P<0.01; ***, P<0.001.

Even though Favipiravir could be efficiently incorporated into RNA products to impair the fidelity of RNA synthesis, it did not significantly inhibit RNA production in vitro (Extended Data Fig. 1). The presence of Favipiravir in RNA product did not immediately terminate the extension of growing strand except when repeat pyrimidine residues were encountered in the template. The RNA synthesis seemed prone to stall if multiple consecutive incorporations of Favipiravir were supposed to take place (Extended Data Fig. 1b), consistent with the observations reported by other groups recently^5^. This evidence suggests that the presence of repetitive Favipiravir residues might distort the configuration of template-product RNA duplex to prevent its further extension. However, the consecutive incorporation of Favipiravir might take place with extremely low probabilities in the cellular environment due to the competition of ATP/GTP substrates. Thus, Favipiravir is more likely to be discretely incorporated during virus replication and induce mutations in progeny RNA copies. This hypothesis was supported by a recently reported virus-based inhibition assay which revealed that Favipiravir was able to escape the proofreading mechanism of SARS-CoV-2 replication complex and led to mutations in progeny viral genome^5^.

To uncover the structural basis of Favipiravir recognition by the polymerase, we determined the structure of SARS-CoV-2 nsp12-nsp7-nsp8 core polymerase complex in the presence of a template-product partial duplex RNA and Favipiravir at 3.2 Å global resolution by cryo-electron microscopy (cryo-EM) reconstruction (Fig. 2; Extended Data Fig. 2). The density map reveals clear features for most amino acid side chains and nucleotide base moieties, allowing the detailed analysis for protein-RNA interactions and Favipiravir recognition (Extended Data Fig. 3). In the structure, the nsp8 subunit (nsp8.1) in the nsp7-nsp8 heterodimer is mostly unresolved, similar to the recently reported structure of Remdesivir-bound complex in the pre-translocation conformation^30^ (Fig. 2a). Besides, the N-terminal long helices of nsp8 subunits could not be visualized as well, which are supposed to form a sliding platform for template-product duplex elongation^27,29^. This might result from the insufficient length of the RNA duplex to establish extensive interactions with the helix track. This structure resolved 18 nucleotide residues in the template strand and 16 residues in the primer strand. Importantly, Favipiravir was captured in the triphosphate form (FTP) before catalysis which bound at the +1 position and paired with a C template residue (Fig. 2b and c). The α-phosphate of FTP is located in vicinity of the 3’-hydroxyl group of A^−1^ residue with a distance of ~3.3 Å and no density for a phosphate-di-ester bond could be observed (Fig. 2c). Moreover, the density of FTP could only be visualized at lower contour levels than the other residues, suggesting the lower occupancy of Favipiravir in the complex, which might be related to the pre-catalytic state without stable covalent interactions with the product RNA. Thus, this structure provides a missing snapshot of the pre-catalytic conformation of SARS-CoV-2 polymerase, an earlier stage before the previously reported post-catalysis/pre-translocation and post-translocation conformations^27,29,30^.

**Fig. 2.**
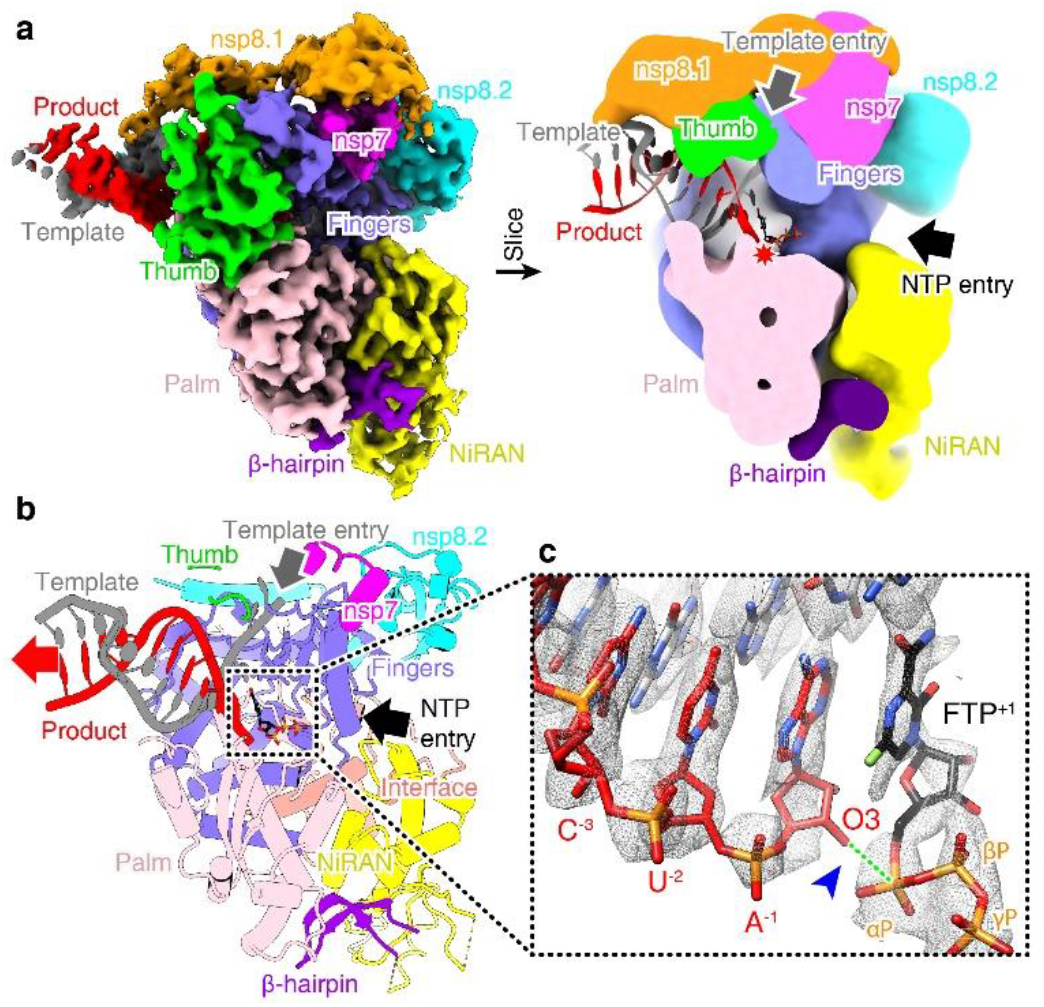
Cryo-EM structure of Favipiravir bound to replicating core polymerase complex of SARS-CoV-2. **a,** Overall density map (left) of SARS-CoV-2 core polymerase complex bound to template and product RNA strands and FTP. The nsp12 polymerase subunit is colored by domains. The nsp7-nsp8 cofactors and RNA strands are colored by chains. The central section (right) of density map is shown to reveal the inner tunnels within the polymerase. The RNA strands are shown in cartoons and colored by chains. The entrances for template and NTP substrate are indicated by arrows. The catalytic site is highlighted by a red star. **b,** Atomic model of the Favipiravir bound polymerase complex of SARS-CoV-2. The protein and RNA are shown in cartoons with the same color code as in **(a)**. The FTP is shown as stick model and highlighted in black. **c,** Close-up view of FTP and the growing terminus of product RNA strand. No density for a covalent bond is observed between O3 of A-1 and α-P of FTP, highlighted by a blue arrowhead. The two atoms are linked by a green dashed line to indicate the position for catalysis.

As a mimicry of nucleotide substrate, FTP is recognized as an incoming NTP by conserved catalytic residues (Fig. 3a). Residues D623 (motif A) and N691 (motif B) stabilize the ribose ring of FTP, and residue S682 (motif B) may potentially interact with both the ribose and pyrazine branch moieties. The A^−1^ residue of product RNA potentially further contact with the α-phosphate and ribose of FTP^+1^ to drag the two residues in close proximity for catalysis. The amide group of Favipiravir pairs with the template C residue and potentially forms three hydrogen bonds for base-pairing (Fig. 3a). When paired with a U template residue, there might be two hydrogen bonds in between (Extended Data Fig. 4). In order to capture the pre-catalytic conformation, we did not include catalytic metals in the buffers for protein purification and assembled the complex by incubation on ice to prevent catalysis (Extended Data Fig. 1c). Yet, we observed the density for a single metal ion anchored by D761 (motif C) of nsp12 which is offset the catalytic position and may arise from the cellular environment during expression (Fig. 3a). A similar conformation was also observed in the structure of enterovirus 71 (EV71) polymerase (PDB: 5F8I)^33^. To enable catalysis, two metal ions are required to neutralize the negative charges of phosphate groups in NTP substrate and stabilize the transition intermediate^33,34^. Based on the previous structure of post-catalytic conformation in the presence of Remdesivir^30^, we modeled two magnesium ions into our structure to analyze the potential catalytic state of this polymerase. The two metal atoms are coordinated by D618 (motif A) and D761 (motif C) and further bridge the three phosphate groups of FTP substrate. The terminal γ-phosphate is further sequestered by a salt bridge contributed by K798 (Fig. 3a). Besides, residues R553 and R555 from motif F may also interact with the β- and γ-phosphate groups. These interactions together stabilize the incoming nucleotide and 3’-terminus of product RNA in close vicinity to facilitate the nucleophilic attack to α-phosphate by the 3’-hydroxyl oxygen of −1 nucleotide.

**Fig. 3.**
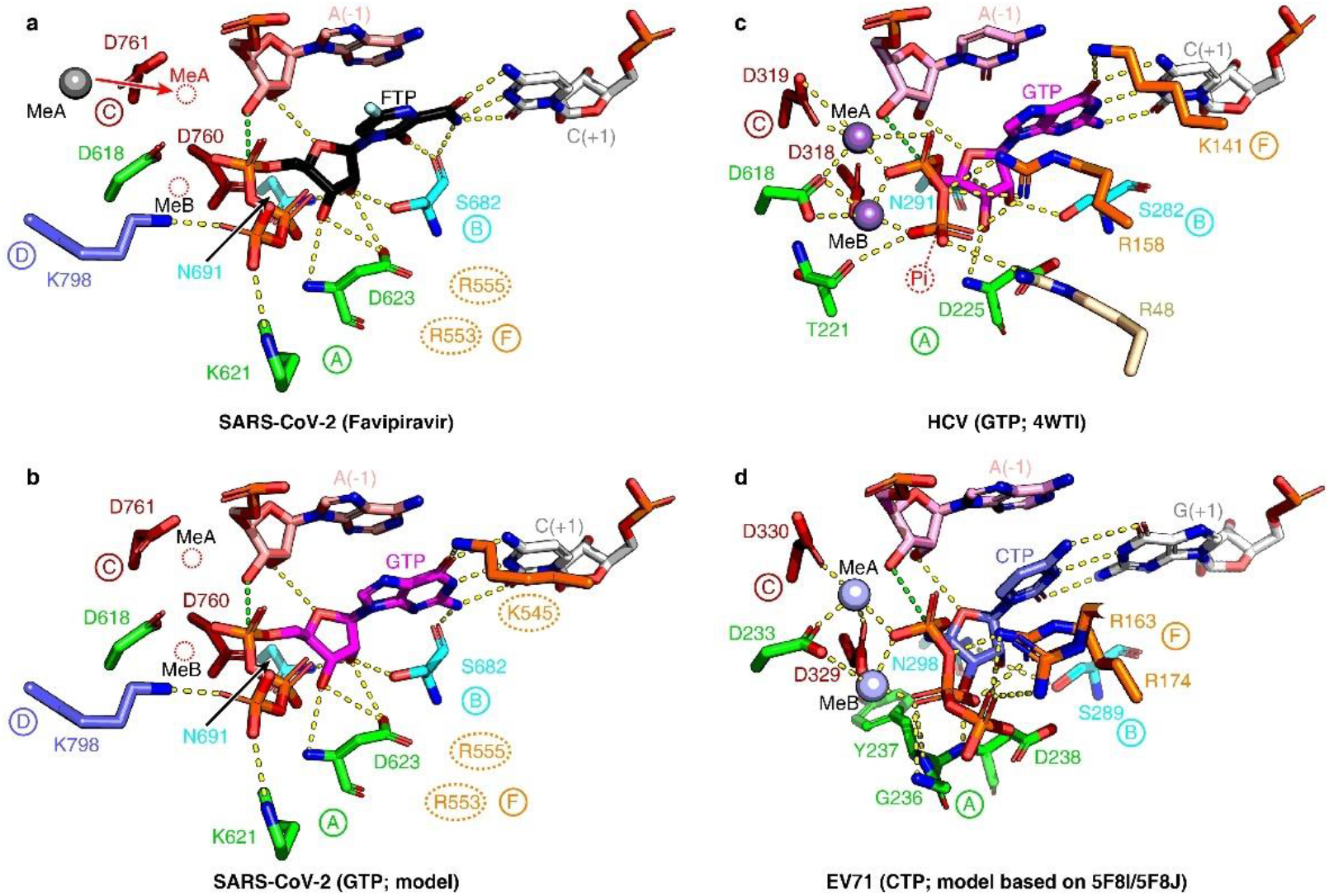
Recognition of Favipiravir and comparison with other NTP substrates. **a,** Recognition of Favipiravir by SARS-CoV-2 polymerase in the pre-catalytic conformation. The key residues of nsp12 involved in FTP interaction are shown as sticks and colored by different catalytic motifs. Positions of the two catalytic metal ions are modeled based on the post-catalytic structure in complex with Remdesivir (PDB: 7BV2)^30^, which are indicated by red dashed circles. The density for a metal ion is observed offset the catalytic site, which presumably corresponds to the catalytic metal A (MeA) before being relocated to the active site, similar to the scenario observed for EV71 polymerase^33^. Two residues in motif F (R553 and R555) are potentially involved in interactions with FTP but are poorly resolved in the density map, represented by dashed ovals. The potential polar contacts are shown as yellow dashed lines. The O3 atom of A^−1^ and the αP atom of FTP is linked by a green dashed line to indicate the site of catalysis. **b,** Predicted model of GTP recognition by SARS-CoV-2 polymerase complex. An additional residue in motif F (K545) is potentially involved in interactions with the guanine base. **c,** Recognition of GTP substrate by HCV polymerase (PDB: 4WTI)^32^. The γ-phosphate of GTP is not present in the structure, indicated by a dashed circle. A residue outside the seven conserved catalytic motifs of RdRp (R48) stabilizes the β-phosphate of GTP by a salt bridge, colored in wheat. **d,** Structural model of CTP bound to enterovirus 71 (EV71) polymerase in the closed conformation. This model was generated based on the pre-catalytic complex in open conformation (5F8I) and the closed post-catalytic conformation (5F8J) of the same complex^33^. All four panels represent the closed conformation in the pre-catalytic state of different structures to facilitate comparison.

Since Favipiravir could mimic both adenine and guanine nucleotides for RNA synthesis, we modeled the GTP into the structure to compare the potential different recognition patterns for different NTP substrates (Fig. 3). Basically, the GTP reveals a highly similar interaction network with the polymerase residues to that for FTP except that an additional residue K545 from motif F may also be involved in interactions with the carbonyl group of the guanine base (Fig. 3b). The ATP substate could also be accommodated similarly with the amino group of adenine base interacting with K545. For pyrimidine nucleotides, however, the base moiety is too far away from K545 to form such interactions (Fig. 3d). This residue is highly conserved in all viral RdRps and may serve as a signature residue for discriminating purine and pyrimidine substrates (Extended Data Fig. 5). Comparing the RdRps from different viruses, including positively-/negatively-sensed and segmented/non-segmented RNA viruses, we found the residues for stabilizing the ribose, base and catalytic metals are highly conserved across all viral families, whereas the residues for accommodating the phosphate groups reveal obvious diversity and may also involve residues outside the seven canonical catalytic motifs of RdRp, e.g. residue R48 in Hepatitis C virus (HCV) polymerase^32^ (Fig. 3c).

With the available structures of SARS-CoV-2 polymerase before and after catalysis^27,29,30^, we were able to assemble a complete scenario of the catalytic cycle to analyze the potential conformational changes of the polymerase during catalysis (Fig. 4). It has been established in flavivirus and enterovirus RdRps that the binding of incoming NTP substrates would induce active site closure and relocation of metal ions to facilitate catalysis^32–34^. A similar scenario may also exist for coronavirus polymerase, and potentially other viral RdRps. The motif A loop in apo polymerase adopts an open conformation with the active site fully accessible for incoming NTP substrates (Fig. 4a and b). One of the catalytic metal ions (metal A) is anchored by D761 offset the catalytic site, whereas the other one (metal B) might be less stable and frequently exchange between the polymerase and the solvent. Therefore, the metal A atom could be visualized in several viral polymerases without the template RNA and NTP substrate in the active site, including bunyavirus (PDB: 6Z6G)^35^, and arenavirus (PDB: 6KLC and 6KLD)^36^ polymerases. With the NTP substrate coming in, the motif A loop flip inward to close the active site and the side chain of D761 (motif C) rotates to relocate metal A into the catalytic site. Besides, residue K798 (motif D) would also be re-oriented to stabilize the γ-phosphate of NTP (Fig. 4c and d). After catalysis, the motif A loop and residue K798 would move back to the open conformation, allowing the pyrophosphoric acid fragment to be released. This process is accompanied by the translocation of template-product RNA duplex, which resumes the polymerase to get ready for the next round of catalysis (Fig. 4e and f).

**Fig. 4.**
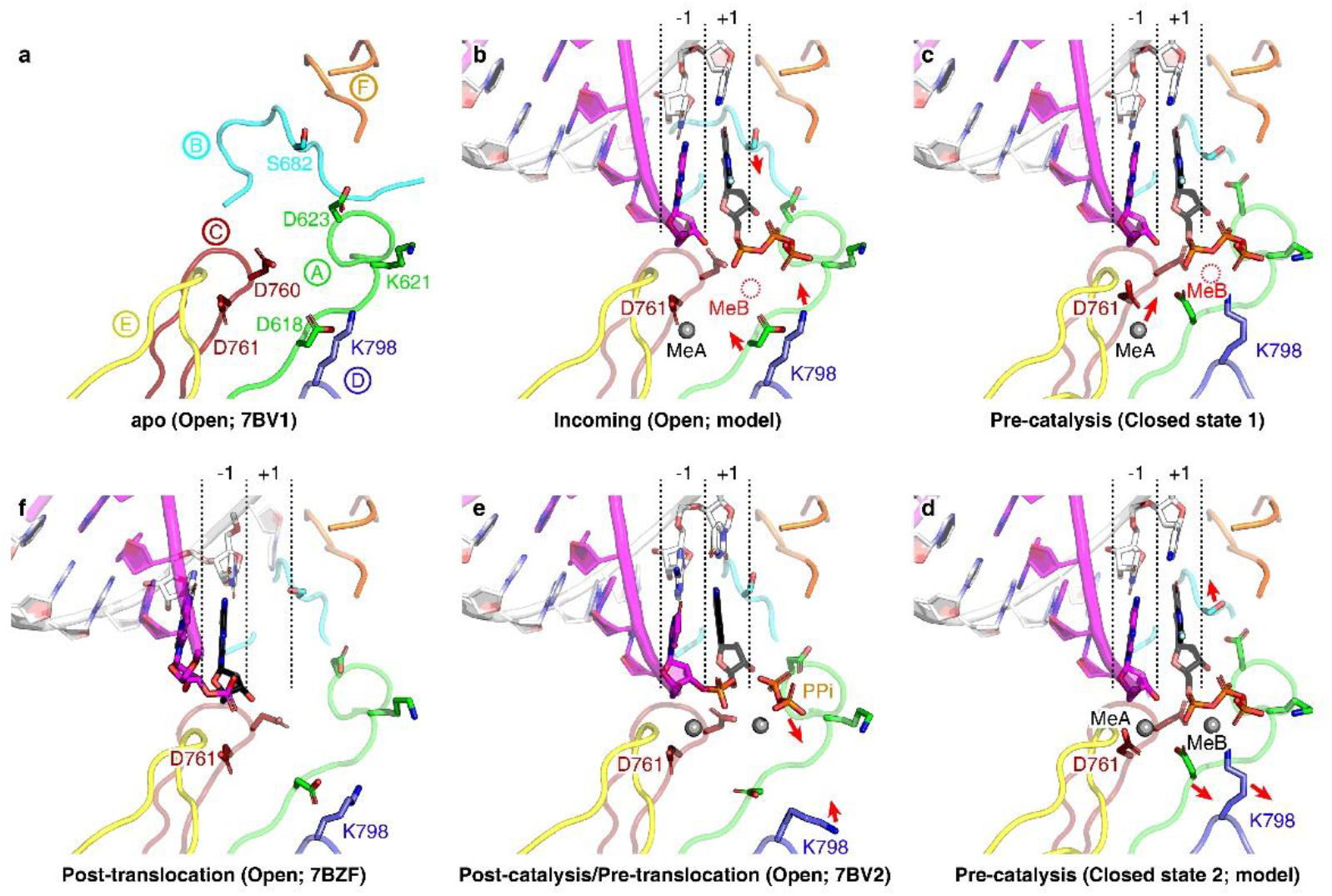
Catalytic cycle of SARS-CoV-2 polymerase. **a,** Catalytic motifs of the apo SARS-CoV-2 polymerase complex. Each motif is represented by a unique color. This structure represents the open conformation of the polymerase. **b,** Predicted model of the SARS-CoV-2 polymerase with NTP substrate coming in. This structure is generated by modelling the RNA and FTP substrate of the pre-catalytic complex into the apo polymerase complex (PDB: 7BV1)^30^. The metal A atom is also modeled offset the catalytic site, which interacts with residue D761. The movements of key residues or motifs for closing the active are indicated by red arrows. **c,** The early-stage pre-catalytic structure in the closed conformation (the reported structure in this study). The metal A is offset the catalytic site and should be relocated as indicated to enable catalysis. The metal B is absent in the structure, indicated by a red dashed circle. **d,** The predicted structure of closed conformation before catalysis. The two metal ions are modeled based on the post-catalytic structure (PDB: 7BV2)^30^. After catalysis, the motifs A and B, and residue K798 would shift to re-open the active site, indicated by red arrows. **e,** The post-catalytic structure of SARS-CoV-2 polymerase in the open conformation before translocation (PDB: 7BV2)^30^. The pyrophosphoric acid fragment would be released and the side chain of K798 would resume as indicated by arrows. **f.** The structure of SARS-CoV-2 polymerase in open conformation after translocation (PDB: 7BZF)^29^.

In summary, we present a missing structural snapshot of coronavirus polymerase replication in the pre-catalytic conformation, facilitating the extrapolation of the dynamic catalytic cycle for RNA nucleotide polymerization. Importantly, we reveal the structural basis of Favipiravir incorporation by SARS-CoV-2, which suggests the feasibility of developing other nucleotide-mimicking antiviral drugs by utilizing non-base derived molecular entities. Given the better record of Favipiravir than Remdesivir in side effect manifestations^3,4^, it may represent a better option for clinical treatment of SARS-CoV-2 infections, as well as a panel of other human-infecting RNA viruses^37^. In addition, these findings provide an important basis for developing better broad-spectrum inhibitors with higher potency to combat the infection of various emerging RNA viruses.

## Methods

### Protein expression and purification

The SARS-CoV-2 nsp7, nsp8 and nsp7L8 fusion proteins were expressed in *E. coli*, and the nsp12 polymerase subunit was expressed with the Bac-to-Bac system (Invitrogen) as previously described^28^. All these proteins were purified by tandem affinity chromatography and size-exclusion chromatography (SEC) accordingly. To constitute the core polymerase complex, the purified nsp12, nsp8 and nsp7L8 proteins were incubated on ice overnight with a molar ratio of nsp12:nsp8:nsp7L8=1:3:3. The complex was then purified by SEC using a Superdex 200 increase column (GE Healthcare) equilibrated with a buffer consisting of 25 mM HEPES-NaOH (pH 7.5), 300 mM NaCl and 2 mM Tris (2-carboxyethyl) phosphine (TCEP). The fractions for nsp12-nsp8-nsp7L8 complex were pooled and concentrated to 4 mg/mL for subsequence experiments.

### In vitro polymerase activity assay

The activity of SARS-CoV-2 polymerase complex was tested as previously described^28^ with slight modifications. Briefly, 40-nt template RNA strands (sequences adapted for each specific NTP substrate as shown in Fig. 1) were annealed to a complementary 20-nt primer containing a 5’-fluorescein label (5’FAM-GUCAUUCUCCUAAGAAGCUA-3’, Takara). To perform the primer extension assay, 1 μM nsp12, 1 μM nsp7 and 2 μM nsp8 were incubated for 30 min at 30 °C with 1 μM annealed RNA and 0.5 mM individual NTP or drug in a reaction buffer containing 10 mM Tris-HCl (pH 8.0), 10 mM KCl, 1 mM TCEP and 2 mM MgCl_2_ (freshly added prior to usage). The products were denatured by heating to 100 °C for 10 min in the presence of formamide and resolved by 20% PAGE containing 9 M urea which was run with 0.5×TBE buffer. Images were recorded with a Vilber Fusion imaging system. The RNA products were quantified by integrating the intensity of each band using ImageJ software. The significance of difference was estimated by the two-tailed student’s t-test to calculate the *P* values for each experimental group.

### Cryo-EM sample preparation and data collection

To prepare the Favipiravir bound polymerase complex, the purified nsp12-nsp8-nsp7L8 complex was mixed with annealed RNA duplex (Template: 5’- CUAUCCCCAUGUGAUUUUACUAGCUUCUUAGGAGAAUGAC-3’, Primer: 5’- GUCAUUCUCCUAAGAAGCUA-3’) with a molar ratio of nsp12:RNA=1:1.5. The mixture was incubated on ice for 2 h in a buffer containing 25 mM HEPES-NaOH (pH 7.5), 150 mM NaCl, 1 mM TCEP and supplemented with 0.5 mM FTP. An aliquot of 3 μL protein solution (0.4 mg/mL) was applied to a glow-discharged Quantifiol 1.2/1.3 holey carbon grid and blotted for 2.5 s in a humidity of 100% before plunge-freezing with an FEI Vitrobot Mark IV. Cryo-samples were screened using an FEI Tecnai TF20 electron microscope and transferred to an FEI Talos Arctica microscope for data collection. The microscope was operated at 200 kV and equipped with a post-column Bioquantum energy filter (Gatan) which was used with a slit width of 20 eV. Cryo-EM data was automatically collected using SerialEM software (http://bio3d.colorado.edu/SerialEM/). Images were recorded with a Gatan K2-summit camera in super-resolution counting mode with a calibrated pixel size of 1.0 Å at the specimen level. Each exposure was performed with a dose rate of 10 e^−^/pixel/s (approximately 10 e^−^/Å^2^/s) and lasted for 6 s, resulting in an accumulative dose of ~60 e^−^/Å^2^ which was fractionated into 30 movie-frames. The final defocus range of the dataset was approximately −1.7 to −3.4 μm.

### Image processing

The movie frames were aligned using MotionCor2 to correct beam-induced motion and anisotropic magnification^38^. Initial contrast transfer function (CTF) values were estimated with CTFFIND4.1 ^39^ at the micrograph level. Particles were automatically picked with RELION-3.0 ^40^ following the standard protocol. In total, approximately 2,895,000 particles were picked from ~5,600 micrographs. After four rounds of 2D classification, ~1,002,000 particles were selected for 3D classification with the density map of SARS-CoV-2 polymerase replicating complex (EMDB-11007) low-pass filtered to 60 Å resolution as the reference. After two rounds of 3D classification, three distinguished 3D classes were identified with clear features of secondary structural elements. These classes showed different extents of flexibility at the distal end of RNA duplex but the main body of polymerase complex was highly similar. Therefore, these 3 classes were combined (including ~329,000 particles) and subjected to 3D refinement supplemented with per-particle CTF refinement and dose-weighting, which led to a reconstruction of 3.2 Å resolution estimated by the gold-standard Fourier shell correlation (FSC) 0.143 cut-off value. In the density map, the long helices within the N-terminus of nsp8 subunits were not observed. To better resolved this region, a local mask for the N-terminal region of nsp8 subunits and the RNA duplex was applied to perform 3D classification without particle alignment. However, no definable class with a long helix track could be identified, suggesting the flexibility of this region in the structure. This phenomenon might result from the shorter RNA duplex observed in other structures^27,29^ that would help to stabilize the long helices. In addition, this structure was restricted by some extent of preferred orientation of particles, which somehow limited the attainable resolution in certain views of the final reconstruction. Basically, the density map was sufficient to support faithful atomic modelling in most regions. The local resolution distribution of the final density map was calculated with ResMap^41^.

### Model building and refinement

The structure of Remdesivir bound SARS-CoV-2 polymerase complex (PDB ID: 7BV2) was rigidly docked into the density map using CHIMERA^42^. The model was manually corrected for local fit in COOT^43^. The structure of FTP was built using the “Ligand builder” plug-in in COOT and manually fitted into the density. The initial model was refined in real space using PHENIX^44^ with the secondary structural restraints and Ramachandran restrains applied. The model was further adjusted and refined iteratively for several rounds aided by the stereochemical quality assessment using MolProbity^45^. The representative density and atomic models are shown in Extended Data Fig. 3. The statistics for image processing and model refinement are summarized in the Extended Data Table 1. Structural figures were prepared by either CHIMERA^42^ or PyMOL (https://pymol.org/).

### Structure-based conservation analysis

The structures of SARS-CoV-2, SARS-CoV (PDB: 6NUR)^46^, EV71 (PDB: 5F8I)^33^, poliovirus (PDB: 3OL8)^34^, HCV (PDB: 4WTI)^32^, Zika virus (PDB: 5TMH)^47^, Vesicular stomatitis virus (PDB: 5A22)^48^, influenza virus (PDB: 4WRT)^49^, Lassa virus (PDB: 6KLC), Machupo virus (PDB: 6KLD)^36^, Severe Fever with Thrombocytopenia Syndrome Virus (SFTSV; PDB: 6L42)^50^, La Crosse virus (LACV; PDB: 6Z6G)^35^ polymerases were aligned with MUSTANG^51^. The aligned sequences were then used for conservation analysis via the ConSurf^52^ server. The figure was generated with PyMOL.

## Data availability

The cryo-EM density map and atomic coordinates have been deposited to the Electron Microscopy Data Bank (EMDB) and the Protein Data Bank (PDB) with the accession codes EMD-30469 and 7CTT, respectively. All other data are available from the authors on reasonable request.

## Acknowledgements

We thank all staff members in the Center of Biological Imaging (CBI), Institute of Biophysics (IBP), Chinese Academy of Sciences (CAS), for assistance with data collection. This study was supported by the Strategic Priority Research Program of CAS (XDB29010000), the National Science and Technology Major Project (2018ZX10101004), National Key Research and Development Program of China (2020YFC0845900), the National Natural Science Foundation of China (NSFC) (82041016, 81871658 and 81802010), and a grant from the Bill & Melinda Gates Foundation. M.W. is supported by the National Science and Technology Major Project (2018ZX09711003) and National Natural Science Foundation of China (NSFC) (81802007). R.P. is supported by the Young Elite Scientist Sponsorship Program (YESS) by China Association for Science and Technology (CAST) (2018QNRC001). Y.S. is also supported by the Excellent Young Scientist Program and from the NSFC (81622031) and the Youth Innovation Promotion Association of CAS (2015078).

## Author contributions

Y.S., Q.P. and R.P. conceived the study. Q.P., B.Y., J.Z. and M.W. purified the protein samples and conducted biochemical studies. Q.P. and R.P. performed cryo-EM analysis. R.P. and J.Q. built the atomic model. Q.P., R.P., M.W., L.F. and Y.S. analyzed the data and wrote the manuscript. All authors participated in the discussion and manuscript editing. Q.P., R.P. and B.Y. contributed equally to this work.

---

**Correspondence and requests for materials** should be addressed to Yi Shi.

**Extended Data Fig. 1.**
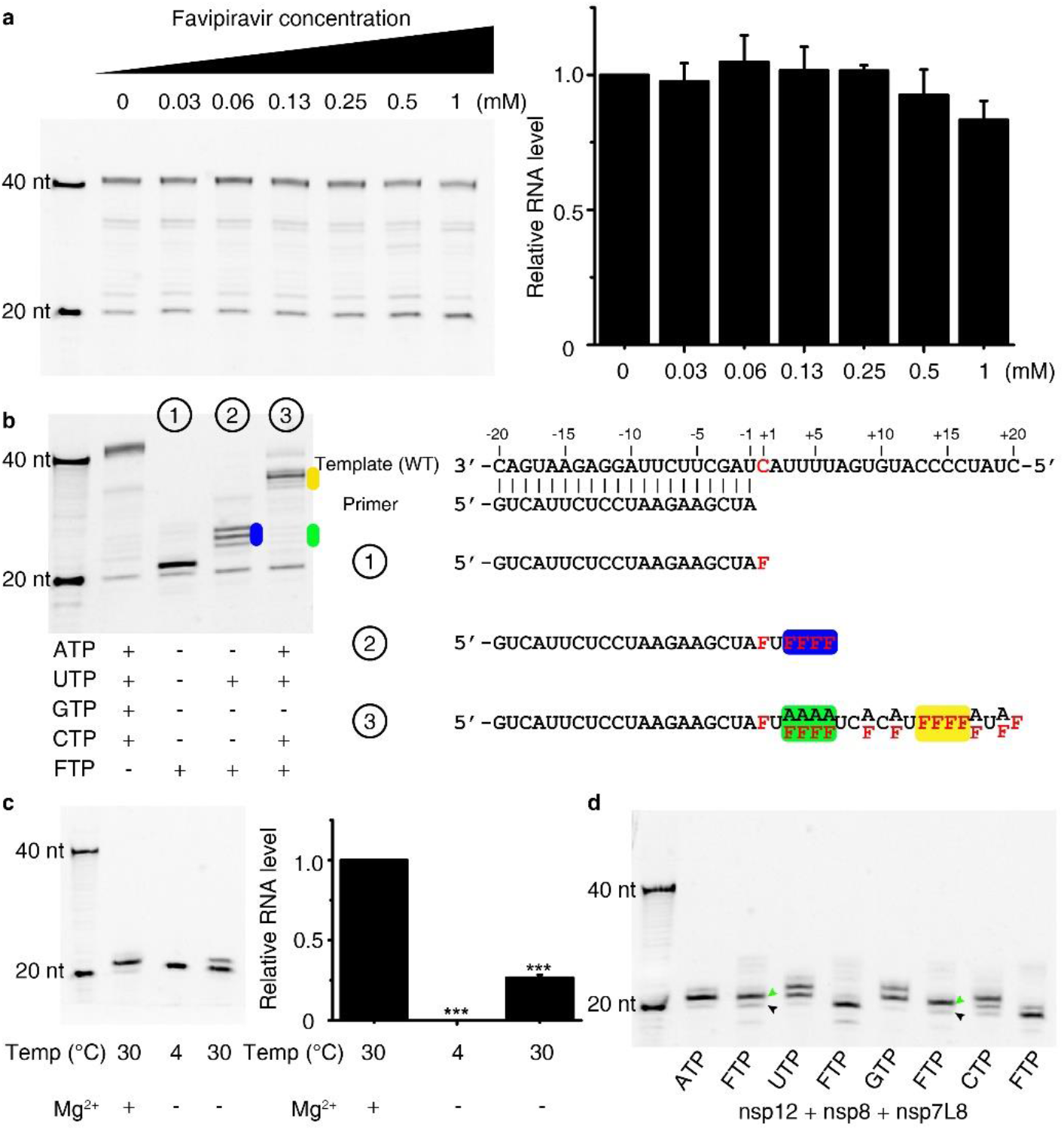
Incorporation of Favipiravir into RNA products by SARS-CoV-2 polymerase. **a,** In vitro inhibition of SARS-CoV-2 polymerase activity by Favipiravir in serially-diluted concentrations. No significant inhibition for RNA synthesis was observed. **b,** RNA extension of in the presence of FTP and different NTP substrates. The RNA synthesis tends to stall at the repeat sequence where consecutive incorporation of FTP is supposed to take place. The potential stalled regions are highlighted in blue, green and yellow backgrounds as indicated in the sequences of RNA products. **c,** The effects of temperature and magnesium ions on the efficiency of Favipiravir incorporation into RNA product. At the ice temperature, almost no catalysis occurred after 2 h incubation in the reaction buffer without magnesium ions. **d,** Catalysis activity of nsp12-nsp8-nsp7L8 complex, related to Fig. 1 for comparison.

**Extended Data Fig. 2.**
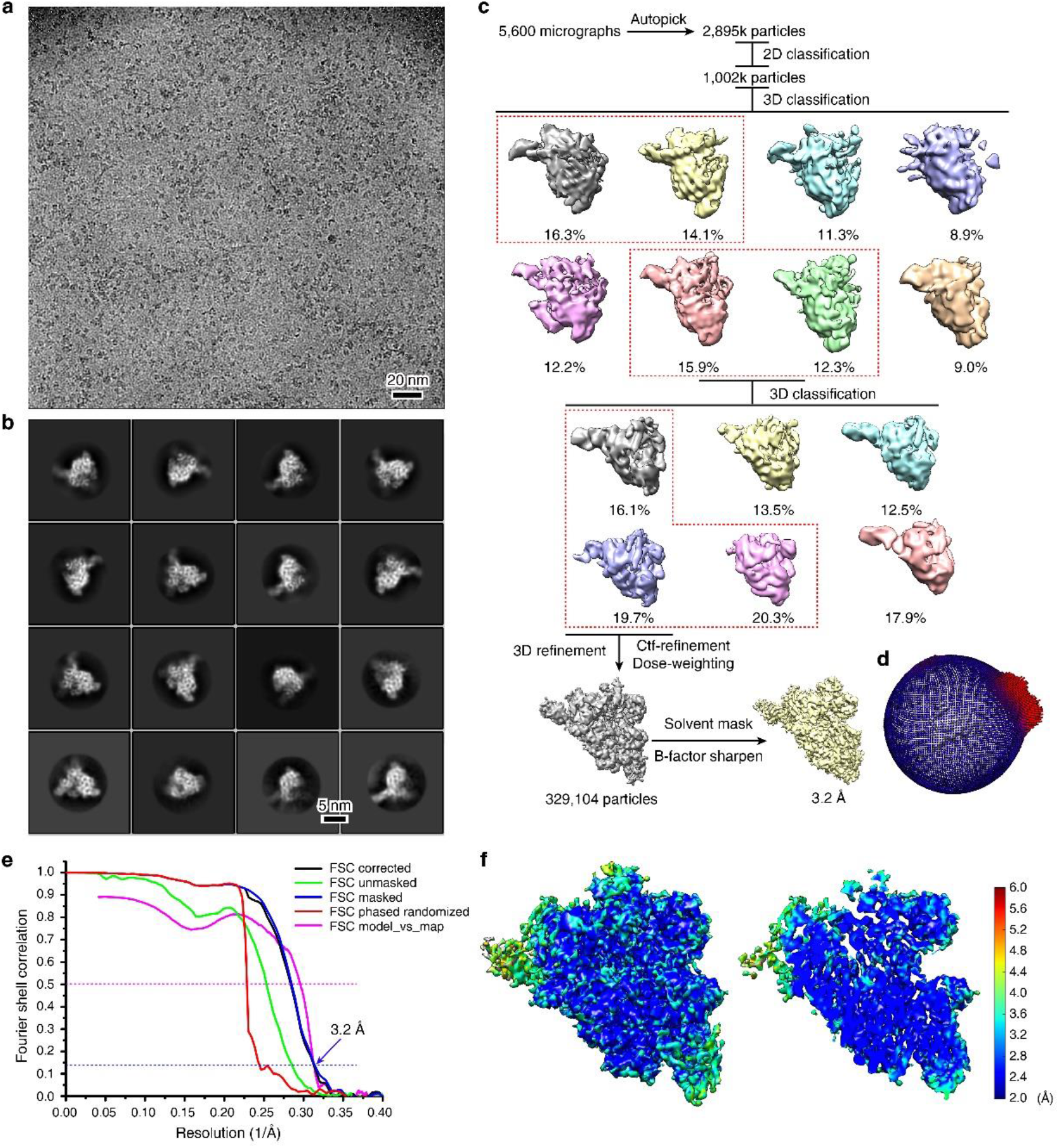
Cryo-EM analysis of SARS-CoV-2 core polymerase complex. **a,** A representative micrograph of SARS-CoV-2 core polymerase bound to RNA and Favipiravir (out of ~5,600 micrographs). **b,** Gallery of 2D class average images. Four rounds of 2D classification were performed. **c,** Flowchart of image processing. The selected classes for next-step processing are indicated by red dashed boxes. **d,** Euler angle distribution of the final reconstruction. **e,** FSC curves for the final reconstruction and model-map correlation. **f,** Local resolution distribution of the final density map.

**Extended Data Fig. 3.**
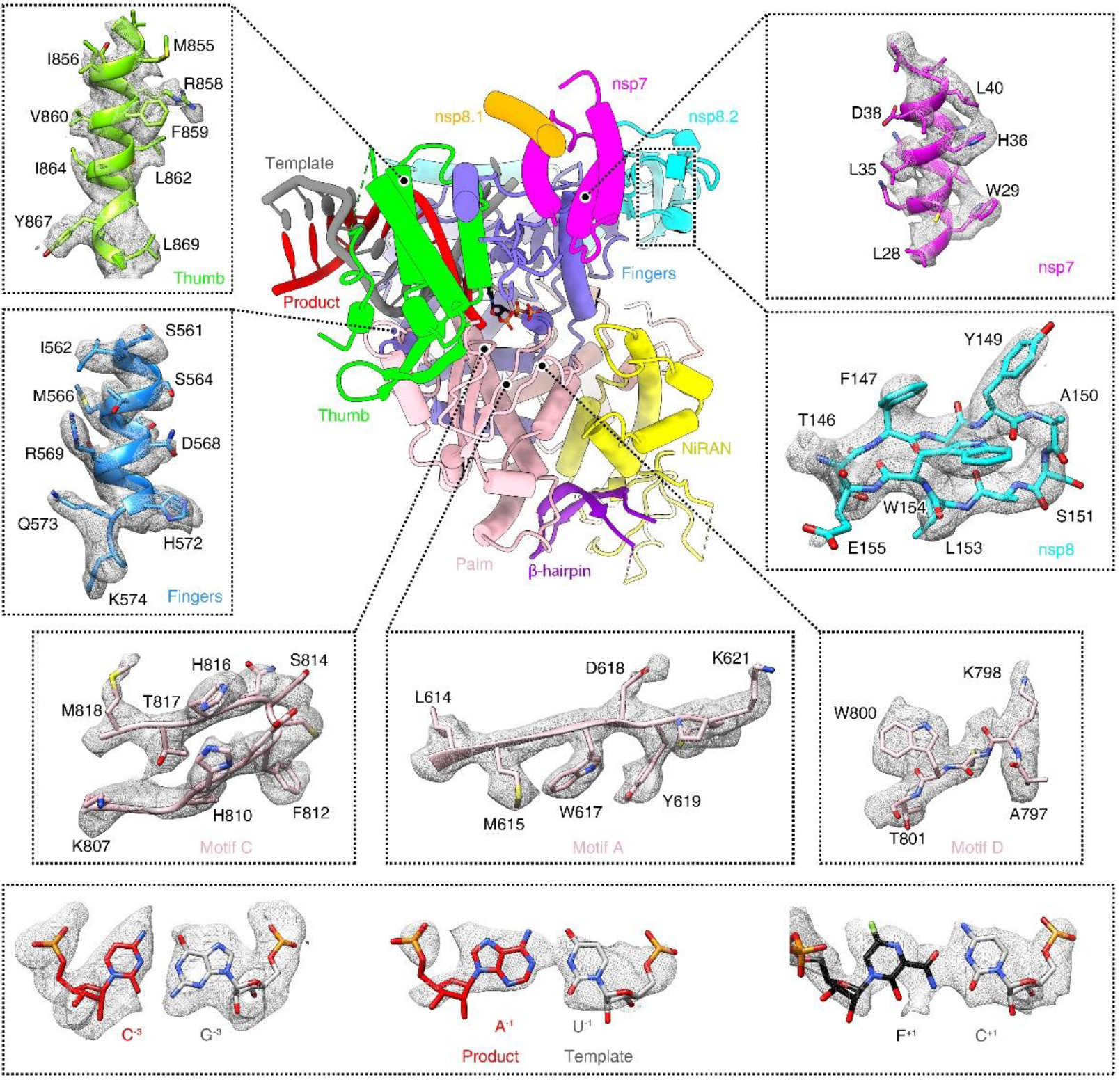
Representative density and atomic models in selected regions. Typical density for different domains, as well as some key residues involved in Favipiravir recognition and catalysis are shown to reveal the quality of the final reconstruction.

**Extended Data Fig. 4.**
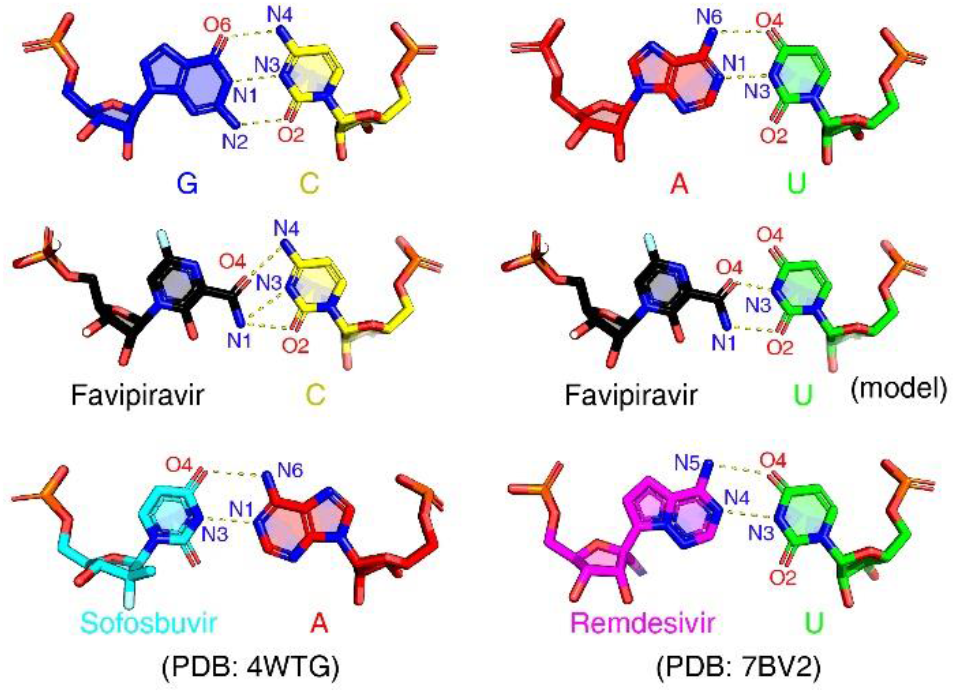
Comparison of hydrogen bond patterns between nucleotide bases and drugs. The interactions within A-U and G-C base-pairs are mimicked by various nucleotide analogue compounds. The hydrogen bonds are represented by yellow dashed lines. The hydrogen donor and acceptor atoms are labeled to facilitate comparison. The structures of Sofosbuvir and Remdesivir are based on their complexes with HCV^32^ and SARS-CoV-2 ^30^ polymerases, respectively.

**Extended Data Fig. 5.**
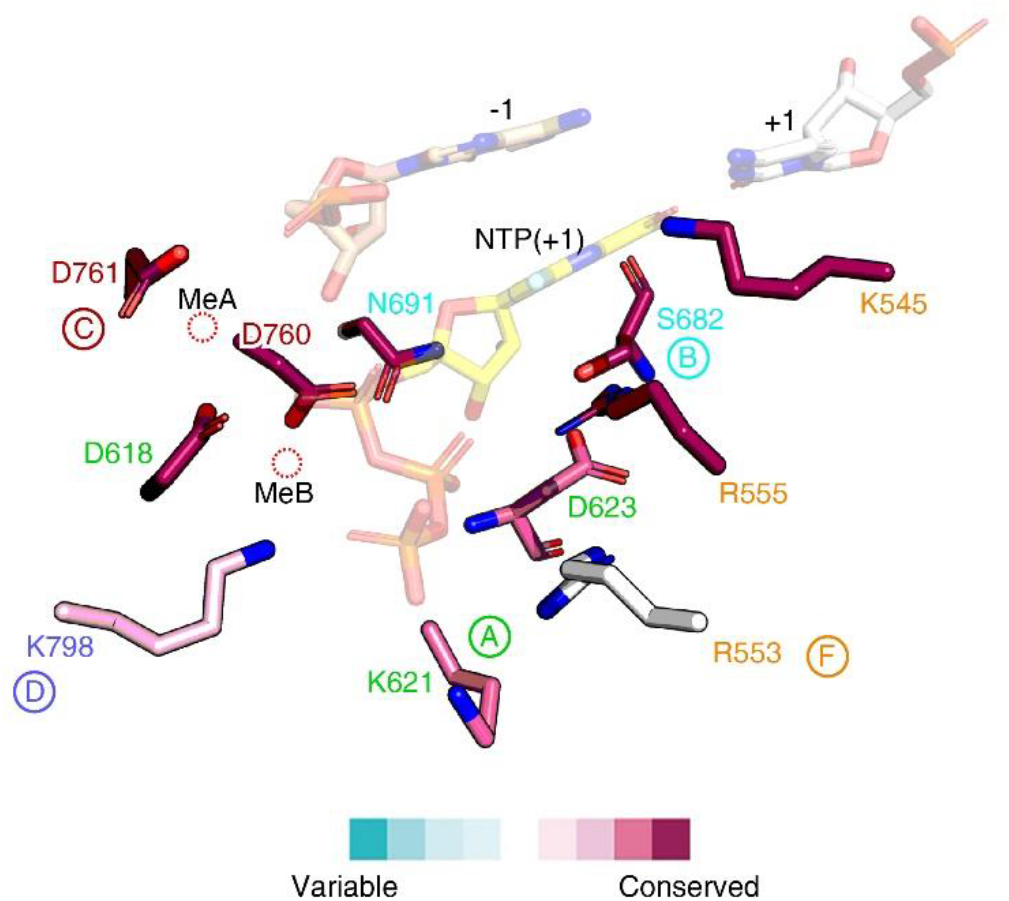
Conservation analysis of key catalytic residues of viral RdRps. The sequences of different viral RdRps are aligned by the available structures in 3D. The conservation scores are then mapped to the structure of SARS-CoV-2 polymerase. The residue labels are colored by the different catalytic motifs as indicated.

**Extended Data Table 1.**
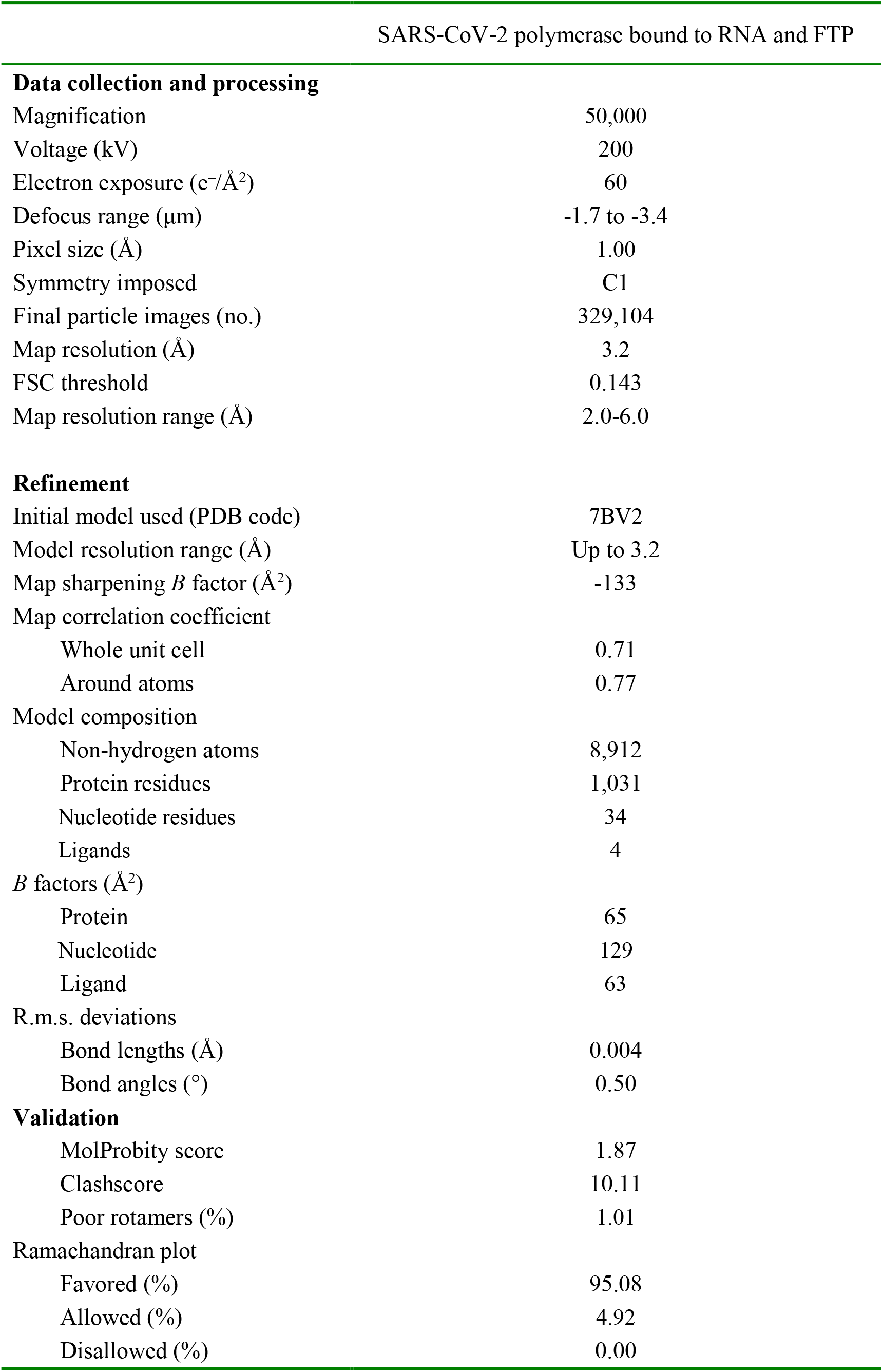
Cryo-EM data processing and refinement statistics.

